# Hippocampal remapping induced by new behavior is mediated by spatial context

**DOI:** 10.1101/2023.02.20.529330

**Authors:** Samuel J. Levy, Michael E. Hasselmo

## Abstract

The hippocampus plays a central role in episodic memory and spatial navigation. Hippocampal neurons form unique representational codes in different spatial environments, which may provide a neural substrate for context that can trigger memory recall or enable performance of context-guided memory tasks. However, new learning often occurs in a familiar location, requiring that location’s representation to be updated without erasing the previously existing memory representations that may be adaptive again in the future. To study how new learning affects a previously acquired spatial memory representation, we trained mice to perform two plus maze tasks across nine days in the sequence Turn Right 1 – Go East – Turn Right 2 (three days each), while we used single-photon calcium imaging to record the activity of hundreds of neurons in dorsal CA1. One cohort of mice performed the entire experiment on the same maze (One-Maze), while the second cohort performed the Go East task on a unique maze (Two-Maze). We hypothesized that CA1 representations in One-Maze mice would exhibit more change in the spatial patterns of neuronal activity on the maze from Turn Right 1 to Turn Right 2 than would be seen in Two-Maze mice. Indeed, changes in single unit activity and in the population code were larger in the One-Maze group. We further show evidence that Two-Maze mice utilize a separate neural representation for each maze environment. Finally, we found that remapping across the two Turn Right epochs did not involve an erasure of the representation for the first Turn Right experience, as many neurons in mice from both groups maintained Turn Right-associated patterns of activity even after performing the Go East rule. These results demonstrate that hippocampal activity patterns remap in response to new learning, that remapping is greater when experiences occur in the same spatial context, and that throughout remapping information from each experience is preserved.

**Significance Statement:** The hippocampus plays a central role in self-localization and the consolidation of new experiences into long term memory. The activity of hippocampal place cells tracks an animal’s spatial location and upcoming navigational decisions, providing, at the ensemble level, unique patterns of activity for experiences that occur in the same physical location. Many studies have demonstrated the existence of divergent patterns at short time scales and how remapping can orthogonalize distinct experiences learned simultaneously. Here, we expand on this knowledge using the power of single-photon calcium imaging to track how new learning affects previously existing spatial memories either in the same or different environments over long periods of time. We observe patterns of hippocampal neural activity in mice during performance of two different rules either in the same environment or in different environments. We find that performing a new behavioral rule in the same environment as a previous rule causes significantly more remapping of hippocampal activity associated with the first rule than observed in mice that perform the two rules in separate environments. However, this remapping does not wholly destabilize memory for the first rule, as many neurons in both groups of mice maintain spatial activity patterns specific to the first rule. These results provide an important step forward in understanding the function of the hippocampus in memory by dramatically expanding the temporal scale over which changes to memory are measured.

## Introduction

A flexible memory system must be able to both retrieve existing memories and rapidly encode new learning. Encoding new learning often involves updating previously existing associations – sometimes with contrasting information – without overwriting previously learned behavioral contingencies because prior knowledge could become adaptive again in the future. Different spatial contexts often dictate distinct behavioral demands, but animals also frequently experience multiple task contingencies in the same spatial environment, requiring them to maintain multiple behavior-context associations and use feedback to adjust their behavior. Playing multiple sports on the same field will lead to associations of two different patterns of behavior in the same physical space: for successful performance, one must use cues beyond the spatial context to discern between memories of multiple behavioral contingencies and follow a single set of rules and skills at any given instance. Additionally, strong memories associated with one sport, such as a crushing defeat, could lead to distraction and affect performance of the other.

The hippocampus enables the use of spatial and behavioral contexts to disambiguate divergent responses to the same stimuli (Brown et al., 2010; Brown & Stern, 2014; Butterly et al., 2012; Komorowski et al., 2009). Populations of hippocampal neurons display orthogonal patterns of activity when spatial environments or behavioral contexts within an environment are sufficiently distinct (Ferbinteanu & Shapiro, 2003; Frank et al., 2000; Griffin et al., 2007; Guise & Shapiro, 2017; Markus et al., 1995; Smith & Mizumori, 2006; Wood et al., 2000). The representational organization of multiple memories is often studied using paradigms in which multiple experiences are learned simultaneously or performed in quick succession (Bladon et al., 2019; Chanales et al., 2017; Guise & Shapiro, 2017; Komorowski et al., 2009; Law et al., 2016; McKenzie et al., 2014; Smith & Mizumori, 2006). Many studies have further shown that representations for concurrently learned, overlapping experiences grow more distinct from each other with continued experience (Brown & Stern, 2014; Chanales et al., 2017; Kinsky et al., 2020; Law et al., 2016; Lever et al., 2002). However, memory representations exhibit continuous change over time via other processes such as sleep, consolidation and representational drift (interchange of neurons participating in a memory ensemble unrelated to changes in environment or behavior) (Cai et al., 2016; Castello-Waldow et al., 2020; Ego-Stengel & Wilson, 2010; Farooq et al., 2019; Girardeau et al., 2009; Rashid et al., 2016; Taxidis et al., 2020; van de Ven et al., 2016). This means that, in behavioral paradigms with short intervals between learning experiences, it is difficult to disentangle the process of encoding and stabilization from the process of orthogonalization. Additionally, the use of short intervals between learned experiences also means that there is little knowledge as to how the process of orthogonalization affects overlapping representations encoded at distant time points.

How existing memory representations accommodate new experience is not well understood. It has been established since the earliest recordings of hippocampal place cells that changes to a spatial environment, such as moving a cue, or to behavioral demands, such as moving a reward location, can elicit remapping of hippocampal activity associated with those features (Lee et al., 2004; McKenzie et al., 2013; Mizuta et al., 2021; O’Keefe, 1976; Shapiro et al., 1997). Similarly, fear conditioning/extinction models of episodic memory have produced evidence that fear learning causes hippocampal remapping (Moita et al., 2003; Moita et al., 2004), and that the process of extinction can both modify the conditioned fear memory association (Nader et al., 2000) and cause encoding of a new, distinct “non-fear” memory trace which suppresses the conditioned fear association (Myers & Davis, 2007). In general, new experiences do not entirely destabilize previously existing hippocampal representations (Cheng & Frank, 2008; Wilson & McNaughton, 1993). Additionally, in familiar environments, a small proportion of place cells can remap their activity to encode new paths (Alvernhe et al., 2011; Lever et al., 2002), reward locations (Boccara et al., 2019; Butler et al., 2019; McKenzie et al., 2013), and fear/threat responses (Moita et al., 2004; Moita et al., 2003; Wang et al., 2012; Zaki et al., 2021). However, in many instances these paradigms do not reinstate a prior behavioral contingency, so in these cases it is challenging to assess whether observed remapping is attributable to representing the new environmental features and behavioral demands, or if it is the result of permanently modifying the previously existing memory representation to reduce interference with new learning.

To study how previously existing spatial memory representations are affected by new learning, we used single-photon miniscopes to image calcium activity of large populations of hippocampal CA1 neurons in two groups of mice performing a sequence of behaviors on a plus maze over nine days. On the first three days, mice followed a Turn Right rule (Turn Right 1); on days four to six, mice switched to a Go East rule; and, for the final three days, mice switched back to the Turn Right rule (Turn Right 2). One group of mice performed the Turn Right and Go East rules on separate mazes (Two-Maze group), and the second group performed the entire sequence on a single maze (One-Maze group). We predicted that forcing the One-Maze group to perform the two different behaviors on the same maze would induce more remapping from Turn Right 1 to Turn Right 2 than would be observed in the Two-Maze group. Additionally, we predicted that Two-Maze mice would form separate contextual representation to disambiguate the rules, which would manifest as a lower correlation of neural activity between Turn Right and Go East sessions than observed in One-Maze animals. Our results are consistent with both hypotheses, but in both groups of mice we also observed a population of neurons which maintained high correlations of activity across the two Turn Right epochs. These data provide physiological evidence for the use of spatial context in the hippocampus to separate representations associated with conflicting behavioral demands and demonstrate that changes to behavioral contingencies alter but do not entirely destabilize hippocampal representations.

## Methods

### Animals and surgery

Six male, naïve mice (C57BL6, Jackson Laboratory) at two-three months of age underwent two stereotaxic surgeries to prepare for calcium imaging, first to infuse virus to transfect neurons with GCaMP6f and second to implant a GRIN lens for imaging. All procedures here were approved by the Institutional Animal Care and Use Committee (IACUC) at Boston University. In both surgeries, mice were given a pre-surgical analgesic of 0.05mL/kg buprenorphine, and were anesthetized with ∼1% isoflurane delivered with oxygen at 1L/min. The first surgery was to infuse virus to express GCaMP6f. During the first surgery, a ∼0.7mm craniotomy was made at AP -2.0mm, ML +1.5mm relative to bregma, above the dorsal hippocampus; then, an infusion needle was lowered at this site to DV -1.5mm relative to bregma, and 350nL of the viral vector AAV9-Syn-GCaMP6f (University of Pennsylvania Vector Core, obtained at a titer of ∼4×10e13GC/mL and diluted to ∼5.5×10e12GC/mL with 0.05M phosphate buffered saline) was infused at 40nL/min and allowed to diffuse for 15 minutes before the infusion needle was slowly removed. The second surgery was performed three weeks later to allow for viral infection and GCaMP6f expression. A 2mm diameter circular craniotomy was made centered at AP -2.25mm, ML +1.8mm, and the neocortex was aspirated while applying near-freezing 0.9% saline solution and GelFoam (Pfizer) continuously to control bleeding until rostral-caudal fiber tracts of the alveus were visible. The GRIN lens (1mm diameter, 4mm length, Inscopix) was slowly lowered stereotaxically to 200 µm dorsal to the infusion site of the virus relative to the skull surface. A non-bioreactive silicone polymer (Kwik-Sil, World Precision Instruments) was used to fix the lens in place, and the polymer and lens were anchored to the skull using Metabond dental cement (Parkell). The lens was covered with removable Kwik-Cast (World Precision Instruments).

After a week of recovery from the lens implantation surgery, mice were again anesthetized and placed in the stereotaxic holder. The miniscope attachment baseplate was placed on the imaging microscope camera, and both were then aligned with the GRIN lens by focusing the entirety of the edge of the lens as viewed in the nVista recording software (Inscopix). The camera with baseplate was then lowered until GCaMP6f-expressing cells were optimally in focus, and then raised by 50 µm to allow for shrinkage of the dental cement used to affix the baseplate. The baseplate was attached to the existing Metabond on the skull with Flow-It ALC Flowable Composite (Pentron), which was then cured with ultraviolet light, and Metabond was applied as an additional layer of reinforcement. In some cases, if clear vasculature were not visible during initial visualization of the brain surface, the GRIN lens was re-covered in Kwik-Cast and the baseplate attachment procedure was performed 1-2 weeks later.

### Imaging

Imaging data were acquired using a commercially available miniaturized head-mounted epifluorescence microscope (Inscopix). Microscopes were attached daily to awake, hand-restrained mice. Optical focus, LED gain and intensity were set individually for each animal. Videos were acquired using the nVista recording software (Inscopix) at 20 Hz with a resolution of 1440 x 1080 pixels, spatially downsampled 2x to 720 x 540 pixels. Dropped and corrupted frames were replaced with the nearest preceding undamaged image, and lost frames were excluded from analysis. Mosaic software (Inscopix) was used to correct for motion of the imaging plane, to crop out large areas without GCaMP6f activity, and create a minimum projection of the final video (the minimum projection of a video is an image made by taking the minimum brightness value of each pixel’s entire time series) to be used during ROI extraction.

TENASPIS software was used to extract neuron regions of interest (ROIs) and calcium event times (Mau et al., 2018; Kinsky et al., 2018; Levy et al 2021) (TENASPIS, software available at https://github.com/SharpWave/TENASPIS; see D.W. Sullivan et al., 2017, Soc. Neurosci. Abstract). In short the TENASPIS algorithm applies a band-pass filter to each imaging video and uses an adaptive thresholding process on a frame-by-frame basis to find connected pixels with discrete patterns of fluorescent activity (blobs). Probable cell blobs are aligned through the imaging time series and determined to be neurons by first computing the cell region of interest’s fluorescence time series, and from there using prior expectations of calcium activity event durations, neuron size and shape, and probable event spatial origin. This algorithm allows for detection of partially overlapping cells and includes steps to isolate calcium events to individual cells. This process is intended to reduce type 1 errors in false calcium transient detection, but results in increased type 2 errors of probable transient event rejection, though this loss is partially overcome by re-examining each neuron ROI’s fluorescence trace to detect missed calcium transients from sharp peaks in the traces. Event localization can ensure that calcium transients from partially overlapping cells are assigned only to the cell with the highest correlation of pixels with those of the shared calcium event.

### Cell Registration

To align neuron ROIs over sessions, we used a combination of CellReg (Sheintuch et al., 2017) with a custom, automated alignment step. We modified the source code for CellReg to optimize RAM use for handling larger volumes of data, primarily by casting arrays into logical entries and eliminating redundant variables; these modifications can be found in our GitHub repository and were tested against the original code to verify they did not impact the registration procedure itself.

Our custom pre-registration step is an automated procedure designed to align imaging field-of-view images from two different sessions based on the relative positions of cell ROIs rather than, as is typically done, based on finding the translation and rotation that produces maximum correlation of the two field-of-view images. Briefly, the relative layout of cell ROIs is described using the Voronoi tessalation/Delaunay triangulation. This diagram is then used to find, for each neuron, all immediately adjacent neurons (Tier 1) and all neurons adjacent to them (Tier 2) (**Supplementary Figure 1A**). We then calculate all the distances from a neuron to its Tier 1&2 adjacent partners, as well as their angles, relative to the vertical orientation (**Supplementary Figure 1C**). This set of Tier 1&2 angles and distances is computed for a reference session and for a session to be registered, and we create a 2-D histogram of the differences between all Tier 1&2 distances and the differences between all Tier 1&2 angles between the two session (**Supplementary Figure 1D**). We threshold the histogram for the difference of distances at 3 microns, and search for the bin in the difference of angles matrix with the most entries. The logic applied here is that if a cell from one session can be registered to another session, then all of the cells around it that will also be registered should have a very small difference of distances from the cell of interest, and the difference of angles should also span a narrow range. By then finding all the pairs of cells with the most Tier 1&2 angle and distance difference entries in the peak bin, we get a set of candidate cells to be registered across the two sessions (**Supplementary Figure 1E,F**). These potential aligned candidate cells are paired to each other across sessions using the minimum of their angle and distance differences and largest number of Tier 1&2 adjacent pairs, and by using only those cells which create matching Voronoi diagrams based on this subset of potential aligned cells. With these potential cross-session “anchors,” a transformation can be computed between sessions to align the whole image (MATLAB function fitgeotrans). The alignment accuracy is then evaluated with CellReg. We found zero cases where this method of alignment was improved by the correlation method in CellReg, assessed by visual inspection and CellReg statistics.

During preparation of this manuscript, it came to our attention that a similar method had been developed by Ma and colleagues (Ma et al., 2016).

### Behavioral Tracking

Animal location within the maze was recorded using a video camera recording at 640 x 480 pixels mounted above the mazes and the CinePlex V2 tracking software (Plexon). Tracking was performed at 30 Hz, and CinePlex provided the TTL pulse to synchronize the imaging data acquisition with the nVista software. Red and green tracking LEDs were attached to the imaging miniscope, and custom software, written in MATLAB, was used to find the average position of the red and green LEDs, which was validated and corrected manually before being linearly interpolated to the imaging time stamps recorded at 20 Hz.

### Maze Description

All experiments were performed in a single experimental testing room that was rich in high-contrast distal cues. Both mazes were set up at the same time in separate locations within the room, so that one could be seen from the other. Mazes were made of wood; the arms of each maze were 1.75 inches wide and 24 inches long, with walls 1 inch tall that mice could easily peer over. Maze 1 was painted yellow, while Maze 2 was surfaced with an off-white rubber-like non-stick material. Mazes were additionally scented with different cleaning solutions, either Nolvasan (Maze 1) or 70% ethanol (Maze 2). Additionally, while mice were on Maze 1, a “summer night crickets” ambient background soundtrack was played through overhead speakers, while on Maze 2 the soundtrack was “ocean waves shoreline.” During the pre-training phase, an additional two mazes (X and Y) were used, both of which had arms 1.75 inches wide and 12 inches long. For learning the Turn Right task, the Maze X was surfaced with lineoleum tile, and the maze for learning the Go East task, Maze Y, was made with white corrugated plastic.

### Behavior

All mice were previously used in a fear conditioning and extinction study between four and ten months of age (Zaki et al., 2021; Orlin et al., in preparation). They were allowed to rest undisturbed in their home cage for a minimum of one week, and then handled for an additional week for 10-15min/day before being trained for this study.

Mice were trained at 11-15 months of age to perform two different rules on plus mazes: a procedural rule, Turn Right, and an allocentric rule, Go East. All mice were initially shaped to perform these rules on a set of two distinct mazes placed at different locations in the same testing room (Mazes X and Y) to ensure that mice reached highly accurate and stable performance of both rules to allow for a similar recording schedule across animals. Animals were initially allowed to wander the mazes freely for 1-2 days, and rewards were then introduced for the rule assigned to each maze (Turn Right on Maze X, Go East on Maze Y). Mice began each trial on the North or South maze arm, with trial order pseudo-randomly determined to limit the number of sequential trials that began on the same arm. Mice were placed on a pedestal between trials. Prior to the start of each trial, the experimenter placed a reward (0.005mL of S30% sucrose solution) in a well in the appropriate arm and pretended to bait the non-rewarded arm; this was always performed in the same order (e.g. reward/pretend reward E before W) so that the mouse could not solve the task by visual example. For the procedural rule, Turn Right, mice were required to go from the South arm to the East arm, or from the North arm to the West arm. For the spatial rule, Go East, mice were required to go to the East arm from either the North or South arms. Mice were initially allowed to freely wander the maze until they found the reward, but as training progressed the mice were removed from the maze and returned to the inter-trial interval pedestal upon making an incorrect arm entry. Arm-entry biases that emerged during training were corrected by repeating the same trial (starting location) several times, and occasionally allowing the animal to correct incorrect arm entries. Training times were similar across mice (10-15 days).

Following the training stage, mice performed the two rules in the same nine-day recording sequence, following the Turn Right rule for days 1-3, Go East for days 4-6, and Turn Right again on days 7-9 (**Figure 1D**). Mice were occasionally allowed to correct error arm entries early on the first day of each rule block (days 1, 4 and 7) to expedite acquisition of the current rule. Correction trials were excluded from analysis. Mice performed the nine-day recording sequence on a set of mazes (mazes 1 and 2) that were distinct from those used during pre-training.

**Figure 1.**
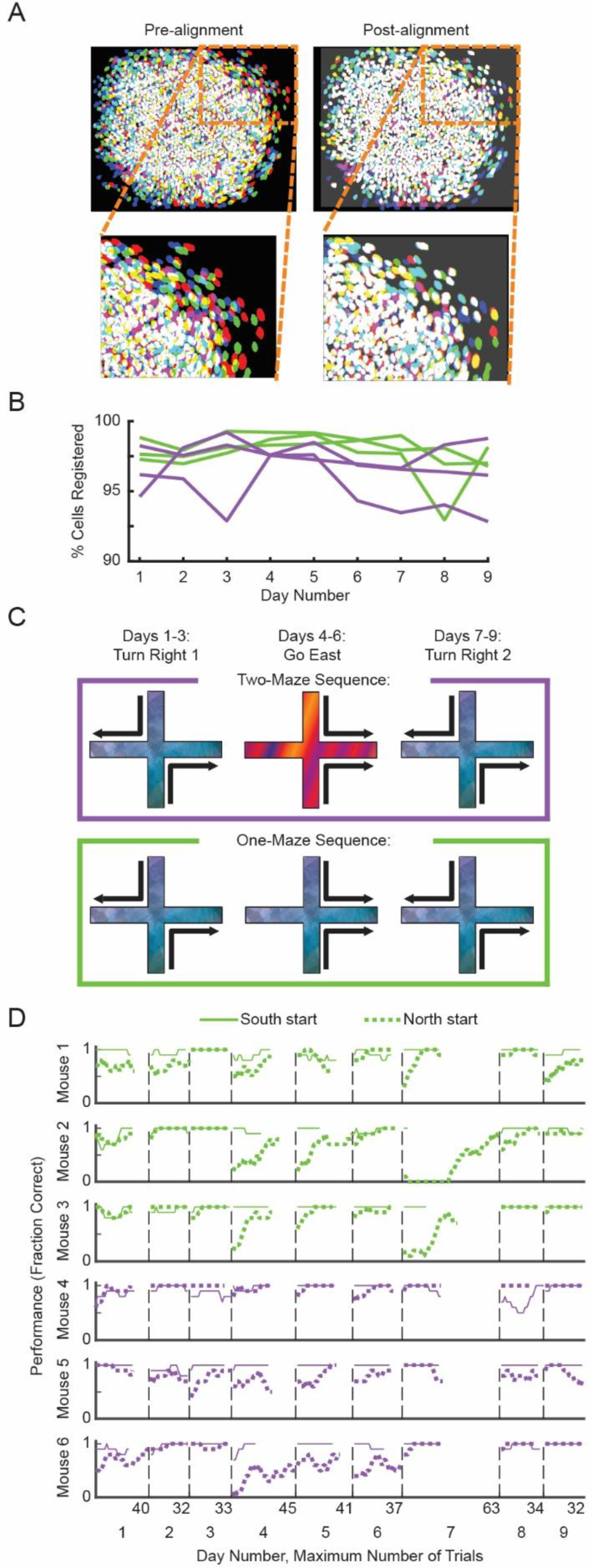
Single-photon calcium imaging in mice performing a nine-day behavioral sequence. **A**, Example neuron regions-of-interest from three recordings days (separate colors) from one mouse, before (left) and after (right) alignment. **B,** The percentage of neurons that were successfully registered on the day indicated to any other recording day. **C,** Schedule of behavioral task and recordings: all mice performed a nine-day sequence alternating Turn Right and Go East tasks in three-day increments. Two-Maze mice performed Go East maze on a second maze, and One-Maze mice performed the whole sequence on a single maze. **D,** Performance of individual mice in a 10-trial sliding window separately for trials starting from North and South. Vertical dashed lines separate recording days, with the maximum number of trials from that day for one start arm indicated at the end of that segment.

### Histology

Mice were sacrificed via transcardial perfusion of 10% phosphate buffered saline followed by 10% phosphate buffered formalin. Extracted brains were post-fixed in formalin for 1-2 days before being transferred to 30% sucrose solution for 2-7 days as a cryoprotectant. Brains were then frozen and sliced into 40 um sections on a Leica CM 3050S cryostat, and sections were then mounted onto slides and coverslipped using Vectashield Hardset mounting medium with DAPI (Vector Laboratories). Images to confirm expression levels of the GCaMP6f protein and location of the virus in CA1 and GRIN lens above were captured using a Nikon Eclipse Ni-E epifluorescence microscope at 10x and 20x magnification.

### Quantification and Statistical Analysis

#### Event likelihood and likelihood rate maps

TENASPIS (see above) computes a local threshold for each neuron ROI’s session averaged fluorescence activity. TENASPIS then returns, for each neuron ROI, a binary series that is true for the rising phase of each significant excursion of the neuron ROI’s fluorescence time series above its local threshold and false otherwise. In this study, calcium event likelihood is calculated by pooling the data from a given set of trials for each spatial bin; counting the number of frames when a given cell was exhibiting a significant calcium event and dividing that number by the total number of frames the animal was in that bin. This produces a value from 0 to 1 describing the likelihood that a neuron was active in that spatial bin.

#### Behavioral restrictions

Neural activity was included only for frames where the animal was moving at least 1cm/second. Trials were only included where the animal made a correct decision on the current task, not including error-correction trials. Activity on the maze was restricted on each arm from 2.86cm to 52.67cm in order to capture the regions where animals’ movements were most consistent to reduce the influence of upcoming turn direction on behavior, exclude the regions where the mice were placed onto the maze each trial, and exclude regions where they received reward. This region was divided into 12 bins, each 4.15cm long.

#### Neuron inclusion activity threshold

In all analyses, neurons were only included if they exhibited a calcium event during at least three laps on one arm. In analyses where arms were considered together (e.g., spatial rate map correlations for all four maze arms across Turn Right 1 and Turn Right 2, or for the three maze arms considered when comparing Turn Right 1 or 2 with Go East), neurons were included if they passed this threshold for at least one maze arm. Across days, we included only neurons that were successfully registered on both days (indicating enough calcium activity to identify an ROI with TENASPIS) and above the activity threshold on at least one day; whether other thresholds were applied (e.g., activity on the maze/arm at all, or being above activity threshold for all days considered) was dependent on the assumptions of each analysis and is indicated in the Results.

#### Modulation Index

The modulation index was adapted from a measure used previously to measure phase locking of gamma activity during the theta rhythm (Tort et al., 2010). First, event counts, in this case trials on which a cell exhibited a calcium event over each arm, are normalized to sum to 1 to get a probability that an event occurs in each arm. We then compute the Shannon entropy of these probabilities, which is -1 times the sum over each arm of the event probabilities, multiplied by the log base 2 of that probability. We then get the KL-distance by taking the log base 2 of the number of arms and subtracting the Shannon entropy. The modulation index value is then calculated by dividing the KL distance by the log base 2 of the number of events.

#### Statistics

Statistical tests were performed using the included MATLAB functions for the Spearman rank correlation, Wilcoxon rank-sum/Mann-Whitney U test, Kolmogorov-Smirnov test, or by custom-written permutation test.

## Results

### Mice Switch Behavioral Contingencies in the Same Environment

Male C57BL6 mice (n=6) performed two behaviors on one or two visually distinct plus mazes over a sequence of days. Mice completed 60-70 trials each day, and trials began on the North or South maze arms following a pseudo-randomly generated order. First, mice were pre-trained on a distinct set of mazes (Mazes X and Y) (Methods) to perform Turn Right and Go East tasks with high accuracy. All mice then performed the Turn Right rule on Maze 1 for three days, (sequence days 1-3, Turn Right 1). Mice quickly reached a high level of performance in the first session and maintained high performance for the next two sessions (**Figure 1D**). All mice then performed the second behavior, Go East, for three days (sequence days 4-6). On the Go East days, mice were assigned to either the One-Maze (n=3) or Two-Maze (n=3) groups: One-Maze mice performed the Go East rule while remaining on Maze 1, and the Two-Maze mice performed the Go East rule on Maze 2. All mice achieved a high level of performance by the third day of Go East (sequence day 6), but One-Maze mice took more trials to achieve high performance. To encourage faster relearning of each rule, we occasionally allowed mice to fix error arm entries or changed the start arm sequence to include more of the trials with high error rates. All mice from both groups then performed Turn Right on Maze 1 for three additional days (sequence days 7-9, Turn Right 2) (**Figure 1C**). All mice re-attained high performance by the third day of Turn Right 2 (sequence day 9) on Maze 1 (**Figure 1D**). All mazes (training and recording) were visually distinct from each other and made of different materials. The two mazes used during the recordings were set up at the same time in the testing room (see Methods).

While mice performed the plus maze tasks, we recorded calcium activity of pyramidal cells expressing GCaMP6f in CA1 of the dorsal hippocampus using single-photon miniaturized microscopes. Cell ROIs and fluorescence traces were extracted from the recordings using TENASPIS (Kinsky et al., 2018; Levy et al., 2021; Mau et al., 2018; see Methods). We found between 662 and 1750 cells per day, and this number was stable within animals over time (within-animal standard deviation across days ∼8.3% of mean number of cells across days). Neurons were registered across days using CellReg (Sheintuch et al., 2017) paired with an additional, custom pre-alignment step (see Methods, **Supplementary Figure 1**) (**Figure 1A,B**). Across all recording sessions from all animals, there were 11,952 unique cell ROIs.

### Greater change in activity from Turn Right 1 to Turn Right 2 in One-Maze Group than Two-Maze Group

In both groups of mice, hippocampal neurons exhibit a wide range of stability and remapping in their spatially tuned patterns of activity over the nine-day recording sequence (**Figure 2A**). To investigate how performing a new behavior in a familiar environment disrupts pre-existing spatial representations, we measured the change in neural activity from the first exposure on the Turn Right task on days 1-3 (Turn Right 1) to the second exposure on the Turn Right task on days 7-9 (Turn Right 2) and assessed differences between the One-Maze and Two-Maze groups. For this analysis, we compared each combination of days from Turn Right 1 and Turn Right 2. For each of these nine pairs of days, we only used cells for which we had a registered ROI on both days in the comparison, had at least one calcium event in the maze region of interest on both days, and passed the activity threshold of being active in at least three laps on one arm for at least one of the days. The arm ends, where the mouse received reward or entered/exited the maze, and the center of the maze were excluded.

**Figure 2.**
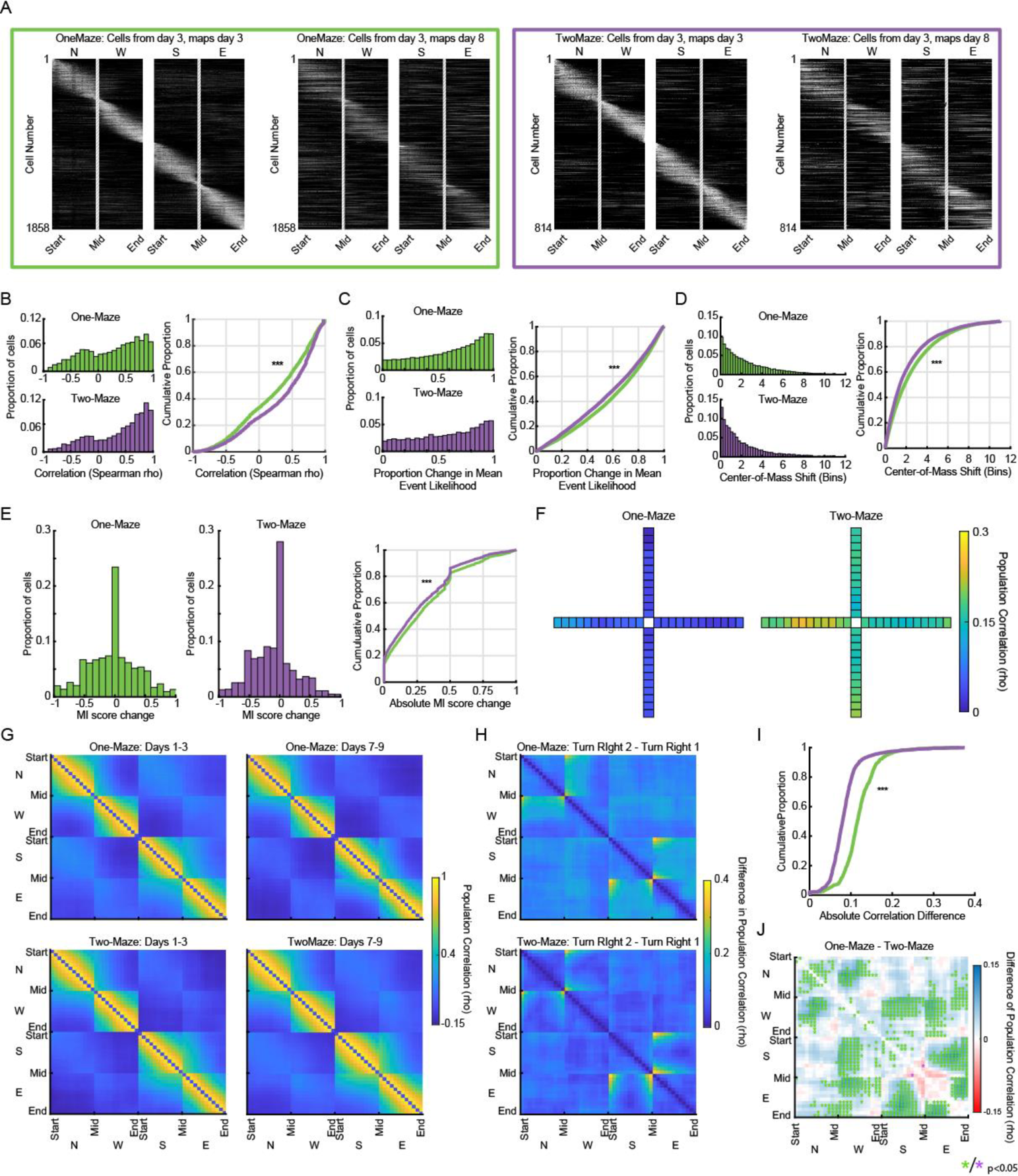
More remapping from Turn Right 1 to Turn Right 1 in One-Maze Mice than in Two-Maze Mice. **A**, Spatial calcium event likelihood maps showing activity from a large number of cells, pooled across animals from each group, from example days 3 and day 8. Each row in a plot is the activity from a single cell, normalized to its own maximum. Rate maps were sorted by maze arm with maximum firing likelihood and center-of-mass of firing within each arm. Rates are normalized to activity on day 3; maps from day 8 are the same cells in the same sort order as on day 3. **B,** Left, histograms for rho values of correlations of single neuron calcium event likelihood maps across all days pairs from Turn Right 1 to Turn Right 2. Top, One-Maze, bottom, Two-Maze. Right, cumulative density plot of same rho values. **C,** Same as B, but for the absolute proportion change of the mean within-arm calcium event likelihood. **D,** Same as B, but for the absolute value of the within-arm shift of the calcium event center-of-mass. **E,** Same as B, but for the change in the modulation index score. **F,** Population vector correlations, averaged across all mice in One-Maze (left) and Two-Maze (right) groups. **G,** Average population vector correlation using activity from within a day, comparing each spatial bin to each other bin; same bin comparisons are set to zero to aid visualization. **H,** Absolute value of differences in within-day bin-to-bin population correlations from G. **I,** Cumulative proportion of correlation differences from H. **J,** Difference of correlation change in each bin pair between One-Maze and Two-Maze activity; blue indicates greater difference from Turn Right 1 to Turn Right 1 in the One-Maze group, red indicates greater difference in the Two-Maze group. Bin-pairs that have an individually significant (p<0.05) difference between groups of animals are indicated with green or purple * for greater difference respectively in One-Maze or Two-Maze groups. *p<0.05, ** p<0.01, *p<0.001

We observed that single neurons in Two-Maze mice had higher calcium event likelihood map (analogous to spiking rate-map; see Methods) correlations from Turn Right 1 to Turn Right 2 than neurons from mice in the One-Maze group (Kolmogorov-Smirnov test, p=6.368e-97, KS stat=0.120; **Figure 2B**). This finding was present separately on all maze arms (**Supplementary Figure 2**). The change of single neuron event map correlations is attributable to changes in both the within-arm calcium event rate (averaged over spatial bins within a maze arm; p=2.922e-19, KS stat=0.053; **Figure 2C**) and location of the within-arm center-of-mass of calcium activity (p=1.993e-58, KS stat=0.093; **Figure 2D**). We additionally computed a Modulation Index (see Methods) which describes how widely activity is distributed across the maze arms: a higher score indicates activity is clustered on fewer maze arms and a lower score indicates activity is spread more evenly across the whole maze. Neurons in the One-Maze group exhibited grater absolute value of change in the modulation index than those in the Two-Maze group (p=6.034e-17, KS stat=0.06, Kolmogorov-Smirnov test; **Figure 2E**). On all metrics, the distributions of remapping scores of from One Maze and Two Maze animals are highly overlapping. These results show that single-neuron remapping induced by performing a new behavior (Go East) in a familiar environment associated with a previous behavior (Turn Right) causes significantly greater change to the hippocampal representation for that previous behavior compared to when individuals perform the new behavior in a separate environment.

To complement the single-unit analyses, we computed population vector correlations to measure changes in ensemble-level activity in each spatial bin. Population vector correlations were significantly higher from Turn Right 1 to Turn Right 2 in the Two-Maze group than in the One-Maze group in all bins except for the three bins at the end of the west arm (all analyses exclude reward location; see Methods) (Wilcoxon ranksum test on group rho values from each day pair; **Figure 2F**). We highlight that this greater level of change in the One-Maze group was observed even on the South and East arms where behavioral demands are unchanged between the two rules.

The results presented thus far indicate that neural activity associated with each arm and spatial bin changed more in the OneMaze group than TwoMaze group, but do not indicate whether such differences extend to the higher-order organization of the representation. State-space population analyses can be used to determine not only how neural populations encode task dimensions, but also how task dimensions are encoded relative to each other, offering a measure of the higher-order structure of a representation. We measured the higher-order structure of hippocampal representations and their change by computing population activity vector correlations for each spatial bin against each other spatial bin within each recording day, and then averaging these bin-by-bin correlations over days 1-3 and 7-9 within experimental groups (**Figure 2G**). Three major observations stand out in the plots of bin-to-bin correlations: first, there was similarity for bins that were close together on the same arm (yellow patches). Second, there were patches of ∼0 correlation between South and East arms and between North and West arms in bins near the middle of the maze (light blue), and also, perhaps unexpectedly, between the two start arms (North and South) and the two reward arms (West and East). And third, the lowest correlations (dark blue) were observed at the start and end arm pairs that would only be paired as mistakes during Turn Right, North-East and South-West. To then examine the magnitude of the change in this structure, we took the absolute value of the difference of these within-day bin-to-bin correlation matrices for each pair of days from Turn Right 1 to Turn Right 2 (**Figure 2H**). Pooling the results from all bin-to-bin comparisons together, there was more change from the One-Maze animals (more yellow colors in top plot of Figure 2H) than in the Two-Maze animals (p=1.591e-151, KS stat=0.499, Kolmogorov-Smirnov test; **Figure 2I**). The difference in the magnitude of bin-to-bin correlation changes was also significantly different in a number of single bin-to-bin comparisons: out of 1128 unique bin pairs, excluding comparison of a spatial bin to itself, there was greater change in One-Maze hippocampal activity differences in 367 bin pairs, and there was greater change in only 1 bin pair in the Two-Maze group (greater change where p<0.05 Wilcoxon rank-sum test for each bin comparison; **Figure 2J**). These analyses show that the combined effects of single-unit remapping also manifest in an altered population code and additionally show that remapping affects not just the representation of spatial bins individually but also the higher-order structure of the representation of the maze environment.

Together, these results demonstrate in a variety of ways that performing a new behavior in an environment previously associated with a different behavior causes substantial change to the hippocampal representation of that environment even when behavioral demands are restored.

### Neurons remap together in coordinated ensembles

The trial-averaged analyses above reveal a bird’s-eye view on the spatially-modulated activity of hippocampal neurons, but additional population activity dynamics are known to exist at much shorter timescales (e.g. Harris et al., 2003). To determine whether additional evidence of activity change related to our task paradigm could be found at the 20-hz sample rate of our recordings of calcium event activity, we made “temporal correlations” for each pair of neurons in a recording session by first concatenating the time series of the calcium event data for each neuron across all trials in a session and then calculating the correlation of these concatenated time series for each pair of neurons. We also constructed “spatial correlations” between pairs of neurons from their session-averaged calcium event likelihood maps over the entire maze (same as in Figure 2). To demonstrate instances where the results of these metrics diverge, we show raster plots of neuronal activity for three cells with well-separated ROIs (**Figure 3A, lower right**): these three cells have high spatial event map correlations to each other (correlations between pairs of cells cell_i_-cell_j_: cells 801-897 rho=0.802, cells 801-956 rho=0.955, cells 897-956 rho=0.854). Two of the cells are preferentially active on later laps and have a positive temporal correlation (cells 801-897 rho=0.409, **Figure 3A top** **row**). In contrast, the third cell (**Figure 3A, bottom left**) is active during early laps in this session and has low temporal correlations with both of the first two cells (cells 801-956 rho=-0.038, cells 897-956 rho=0.020). This example demonstrates that pairs of neurons may be active independently of each other despite having similar place field locations.

**Figure 3.**
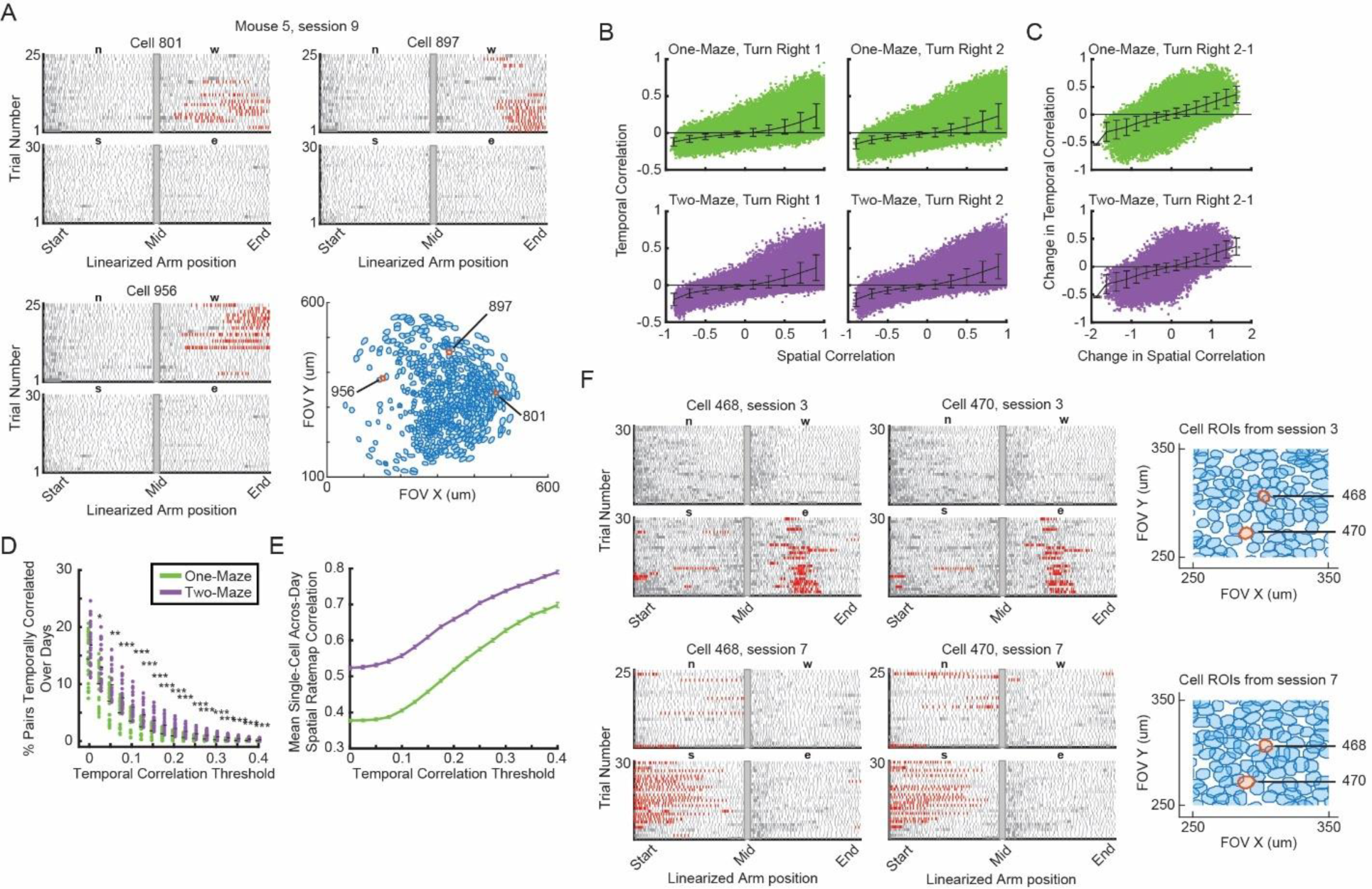
Greater preservation of frame-by-frame activity correlations across neurons in Two-Maze mice than in One-Maze mice. **A**, Raster plots for three example cells from the same recording session, and ROIs for all cells in the same session with the three example cells highlighted in red. In the raster plots, each row of ticks is a single trial, progression from the trial start on the north or south arm to the trial end on the west or east arm. Each tick is the animal’s linearized position on the indicated arm for each frame of imaging. Tick marks are colored pink when that neuron exhibited a calcium transient. **B**, Temporal correlation and spatial correlation of each pair of active neurons from each day of Turn Right 1 (left) and Turn Right 2 (right). Trend line shows mean+/-standard deviation of that decile. **C**, Change in temporal and spatial correlation for pairs of neurons that stay active from Turn Right 1 to Turn Right 2. Trend line shows mean+/-standard deviation of that decile. **D**, Percentage of neurons that stayed correlated over each day pair from Turn Right 1 to Turn Right 2. Correlation threshold indicated on x-axis. Statistic: Wilcoxon ranksum test. **E**, Mean correlation of within-neuron spatial activity maps over days for each neuron that was a member of a pair temporally correlated above the threshold indicated on the x-axis for both days tested. Line indicates mean, error bar standard error of the mean. **F**, A pair of neurons from Mouse 1 that remain temporally correlated and remap together, and a section of the ROI plots from the session days. *p<0.05, **p<0.01, ***<p<0.001

To assess whether the relative time of activation could present another dimension of remapping, we first quantified the divergence in spatial and temporal correlations among pairs of neurons. Since many pairs of hippocampal neurons have overlapping place fields, it is expected that many pairs of neurons in this analysis would exhibit high spatial and temporal correlations; pairs of neurons with non-overlapping place fields will by definition have low temporal correlations. As expected, there was a significant positive relationship between the temporal and spatial correlations from pairs of neurons in both One-Maze and Two-Maze mice during Turn Right 1 and Turn Right 2 (One-Maze, Turn Right 1: rho = 0.650, Turn Right 2: rho=0.675, Two-Maze, Turn Right 1: rho=0.722, Turn Right 2: rho=0.734; all p<0.001; Spearman rank correlation) (**Figure 3B**).

We next assessed the variability in the persistence of spatial or temporal coordination among pairs of neurons over days. First, we asked whether changes in these metrics were coordinated from Turn Right 1 to Turn Right 2, which would be indicated by a positive relationship between the change in spatial correlation and the change in temporal correlation across a pair of days here we found a statistically significant positive correlation in both groups of mice (One-Maze, rho=0.6060; Two-Maze rho=0.6056, both p<0.001) (**Figure 3C**). It is possible that changes in spatial and temporal activation would not necessarily be linked: for example, neurons with similar place fields could independently change which set of laps within a session they are active. Instead, these results suggest that change in when a cell is active relative to another cell is related to change in where the two cells are active.

The above results indicate a preserved relationship between spatial and temporal correlations among all cells that remained active from Turn Right 1 to Turn Right 2, so we next asked whether there were differences between the One-Maze and Two-Maze groups in patterns of activity among cells that remained temporally coordinated. We found that a higher percentage of neurons remained a member of a temporally coordinated pair from Turn Right 1 to Turn Right 2 in the Two-Maze group than in the One-Maze group at most thresholds tested (p<0.05 at threshold 0.025, p<0.01 at threshold 0.05, and p<0.001 all thresholds above 0.075; Wilcoxon ranksum test) (**Figure 3D**). Additionally, the spatial correlation across days was higher in temporally coordinated neuron pairs in the Two-Maze than in the One-Maze group at all temporal correlation thresholds (all p<3.240e-22, Wilcoxon ranksum test) (**Figure 3E**). These results show that temporal coordination among neuron pairs is yet another dimension of remapping that is influenced by performing the Go East task in the same maze as the Turn Right task. Changing temporal correlation is partly independent of other remapping metrics presented in this study, as **Figure 3E** shows that many pairs of neurons remap their place fields together while remaining temporally coordinated (example raster plots shown in **Figure 3F**).

These results show that the greater remapping observed in the One-Maze group from Turn Right 1 to Turn Right 2 was not a random and undirected reorganization of network activity. While neurons in the One-Maze group were more likely to remap across days than neurons from the Two-Maze mice, pairwise neuron coordination in the One-Maze group was preserved as well, just to a lesser degree. Additionally, spatial and temporal correlations are related to each other but are also separable: while neurons that remain members of a temporally coordinated pair over days tend to be more stable, many pairs of neurons also display coordinated remapping by changing their place fields in sync while remaining temporally coordinated. The increase in across-day spatial correlation for members of a highly temporally correlated neuron pairs (**Figure 3E**) could occur through Hebbian-type plasticity mechanisms reinforced by temporal coordination, meaning that temporal coordination can provide a foundation for pairs of neurons to become even more coordinated with each other in time throughout a session, in space across the maze, or both.

### Activity in Two-Maze mice is primarily separated by maze environments, in One-Maze mice by temporal interval

The above results show less remapping of hippocampal neuron activity patterns from Turn Right 1 to Turn Right 2 in the Two-Maze mice than in the One-Maze mice, but they do not address whether remapping associated with transitions to the Go East rule and back to the Turn Right rule involves modification of the existing map or establishment of a new one. Changes in activity from Turn Right 1 to Turn Right 2 may have been induced by both the changes in behavior and by the time between recording sessions (Guise & Shapiro, 2017; Malagon-Vina et al., 2018; Mau et al., 2018).

We first probed whether mice utilized separate neural activity maps to represent the two rules. To compare activity across sessions with different rules, these analyses include all trials on the North and South arms, as well as trials on the East arm that began on the South arm. We first examined the mean population vector correlation for each adjacent day pair (1-2, 2-3, etc.) to look for evidence of changes related to the rule switch. We observed a reduction of the day-to-day population activity correlation in the Two-Maze group on day pairs 3-4 and 6-7, coinciding with transitions between the task rules and maze environments; the activity correlation in the One-Maze group did not visibly decrease over those same day pairs (**Figure 4A**). We next examined the single-neuron calcium event likelihood map correlations over these same pairs of days. In both One-Maze and Two-Maze groups, we observed lower correlations for across-rule day pairs when mice shifted rules than for the other within-rule day pairs (Kolmogorov-Smirnov test, One-Maze p=1.48e-5, KS stat=0.048; Two-Maze p=1.389e-166, KS stat=0.400; **Figure 4B**, upper panels). Notably, the magnitude of this difference was greater in the Two-Maze group, indicating combinatorial effects of the changes in spatial environment and behavioral demand. Overall, there was a difference between groups for both within-rule and across-rule correlations, but this difference was of a smaller magnitude for within-rule correlations (p=1.128e-33, KS stat=0.108) than it was for across-rule correlations (p=9.886e-86, KS stat=0.3001) (**Figure 4B**, bottom panels, same data as top panels).

**Figure 4.**
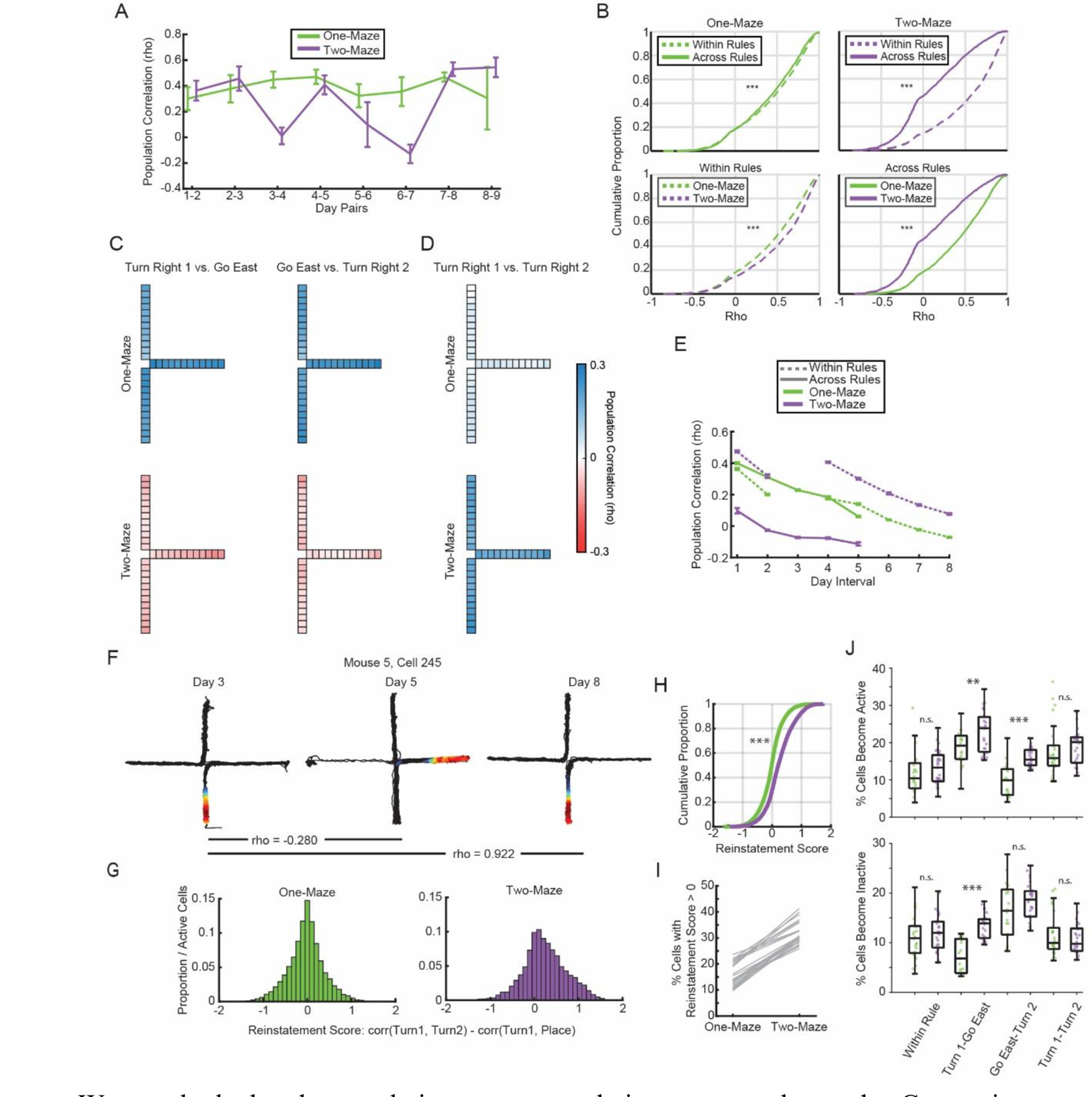
One-Maze neural activity changes slowly over time, Two-Maze neural activity changes sharply with rule and environment. **A**, Population vector correlations for adjacent day pairs. **B**, Cumulative distribution of single neuron event likelihood correlations for day pairs across rules changes against day pairs within rule epochs. **C**, Population vector correlations between Turn Right 1 and Go East day pairs (left), Go East and Turn Right 2 day pairs (right), and **D**, Turn Right 1 and Turn Right 2 day pairs. C and D use the same color scale (right). **E**, Population vector correlations averaged across spatial bins and pairs of days that span the same temporal interval. Top, all spatial bins and temporal intervals grouped together. Bottom, same as top, but separated by whether the days in the pair are from the same or different task rules. **F**, Example firing rate map demonstrating reinstatement of Turn-Right associated pattern of activity during Turn Right 2. Black line is the animal’s trajectory, colored dots are locations where this cell was exhibiting a calcium transient, dots are colored according to the calcium event likelihood within a 5cm radius around each dot, and colors are normalized to the maximum for each session. **G**, Histogram of the differences of correlations for each group of mice. **H**, Cumulative proportion of differences in correlation values across day triplets from Turn Right 1 to Go East to Turn Right 2. One-Maze in green, Two-Maze in purple. **I**, Percentage of neurons with a positive correlation from Turn Right 1 to Turn Right 2 and a negative correlation from Turn Right 1 to Go East. **J**, Percentage of neurons that become active (top) or inactive (bottom) across a day pair for the comparison listed below. *p<0.05, **p<0.01, ***p<0.001

We next looked at the population vector correlations across rule epochs. Comparisons from Turn Right 1 to Turn Right 2 are the same as observed in **Figure 2F**, excluding the West arm (**Figure 4D**), and correlations are higher in the Two-Maze group than in the One-Maze group. From Turn Right 1 to Go East and from Go East to Turn Right 2, neural activity was negatively correlated in Two-Maze mice (**Figure 4C**, bottom) but correlations were relatively high in One-Maze mice (**Figure 4C**, top).

Many previous studies have reported that spatial activity patterns of a neural population continuously change over time due to spontaneous remapping and the exchange of the active population of neurons, a phenomenon known as drift (Kinsky et al., 2020; Levy et al., 2021; Mankin et al., 2012; Mau et al., 2018; Rubin et al., 2015; Ziv et al., 2013). To compare how the change in rule and passage of time each influenced remapping, we plotted the population vector correlation against the interval between recording sessions separately for all across across-rule and within-rule day pairs for each group of animals. We observed drift in our data, as indicated by a decrease in the population correlation with increasing interval between sessions, similar to what has been previously observed (**Figure 4E**). In the One-Maze group, the population correlation monotonically decreases as a function of increasing session interval, and there is a small separation between the lines for session pairs within or across task rules, consistent with what we observed in **Figure 4A,B**. Neural activity in the Two-Maze group also exhibits a time-related decrease in the population correlation, but the effect of performing the Go East rule on a separate maze caused such a large decorrelation of activity patterns that Two-Maze mice exhibited lower correlations across rules at short lags than for pairs of days with the same rules even at longer lags. This result expands on previous findings by showing that patterns of hippocampal neural activity diverge more across different maze environments than for both different task rules and the time between exposures together.

To assess whether the remapping in the One-Maze group represented a complete overwriting of the Turn Right 1 representation by the Go East activity, we next looked for evidence of reinstatement of Turn Right 1-associated activity during Turn Right 2. We looked at single neuron calcium event map correlations for all 27 triplet combinations of days using one day from each of the three rule epochs (Turn Right 1, Go East, Turn Right 2). For each day triplet, we made correlations of the single unit rate maps from Turn Right 1 to Go East and from Turn Right 1 to Turn Right 2. We then calculated a reinstatement score by subtracting the Turn Right 1-Go East correlation from the Turn Right 1-Turn Right 2 correlation: a positive reinstatement score indicates higher a correlation between the two Turn Right epochs, while a negative score indicates a higher correlation from Turn Right 1 to Go East. **Figure 4F** shows example event rate maps from a neuron with a higher activity map correlation between a Turn Right 1 day and a Turn Right 2 day than from the same Turn Right 1 day to a Go East day, and thus a positive reinstatement score for this day triplet. Single neuron reinstatement scores were higher in the Two Maze group than in the One-Maze group (Kolmogorov-Smirnov test, p=0, KS stat=0.263, **Figure 4G,H**). Positive reinstatement scores suggest retrieval of the original calcium event map from Turn Right 1 during Turn Right 2: we additionally found that the Two-Maze group had a higher proportion of neurons displaying “true” reinstatement than did the One-Maze group, meaning these neurons had positive reinstatement scores, positive event map correlations from Turn Right 1 to Turn Right 2, and negative correlations from Turn Right 1 to Go East (15.47±4.34% One-Maze vs. 31.41±4.22% Two-Maze, mean±standard deviation, p=5.606e-06 Wilcoxon signed-rank test, **Figure 4I**; each line corresponds to a day triplet). The presence of a large number of neurons displaying Turn-Right reinstatement in the One-Maze group is notable in that it demonstrates that remapping from Turn Right 1 to Turn Right 2 is a heterogeneous process that partially preserves the hippocampal representation of the Turn Right 1 experience, a result consistent with findings from studies on fear extinction that emphasize inhibition of fear memories rather than their erasure (Bouton et al., 2021).

One method by which a spatial representation could be shielded from interference from a second context is by using a separate population of cells to represent each context. To assess whether this dynamic was present in our data alongside the remapping already described, we looked at whether cells became active or inactive across a pair of days, including only cells for which we could find an ROI on both days of a given pair. Many of these comparisons are consistent with our previous findings and show larger effects in the Two-Maze group than in the one maze group. First, there was no difference between groups of mice in the percentage of neurons that became active or inactive across a pair of days within a rule epoch (become active: p=0.160; become inactive: p=0.456; Wilcoxon ranksum test) or from Turn Right 1 to Turn Right 2 (become active: p=0.207; become inactive: p=0.665)(**Figure 4J**). Next, neurons in Two-Maze mice were more likely than those in One-Maze mice to both become active (p=0.008) and inactive (p=2.2924e-6) from Turn Right 1 to Go East, and from Go East to Turn Right 2 were more likely to become active (p=3.567e-6) but not become inactive (p=0.253). Interestingly, while in both groups more neurons became active over day pairs from Turn Right 1 to Go East than across day pairs within a rule epoch (One-Maze: p=1e-4; Two-Maze: p=5.434e-7), the percentage of neurons that became inactive was greater in the Two-Maze group (p=0.021) and substantially lower in the One-Maze group (p=0.005). Finally, in both groups we saw greater inactivation from Go East to Turn Right 2 than for day pairs within a rule epoch (One-Maze: p=0.003; Two-Maze: p=1.300e-6), but only neurons from the Two-Maze group were more likely to become active in this same comparison (Within Epoch vs. Go East-Turn Right 2: One-Maze: p=0.207; Two-Maze: p=6.144e-4). Overall, these results show that neurons from the Two-Maze mice are more likely than in the One-Maze mice to cross above and below the activity threshold over Turn Right to Go East day pairs, meaning that rather than remap these neurons preferentially fire for a single maze environment.

The results in this set of analyses show that both groups had a mixture of cells that represented each rule type across sessions, but Two-Maze mice tended to better preserve their representations of the Turn Right 1 experience. They further suggest that the reason the Two-Maze mice exhibited less change in the hippocampal representation for the Turn-Right maze and behavior is that they used a separate contextual representation for the maze where they performed Go East, thus reducing interference. In the One-Maze mice, on the other hand the separation of neural activity between the two behaviors is much lower and is closer in magnitude to what is observed as day-to-day drift; in One-Maze mice, drift has a larger effect in decorrelating representations than performing the same behavior in Turn Right 1 and Turn Right 2 has in reinstating the original activity map.

## Discussion

The results of this study show the hippocampal activity code changes more when a second behavior is performed in an environment previously associated with a different behavior, compared to when that second behavior is performed in a separate environment. We recorded hippocampal calcium activity while mice performed two rules in the sequence Turn Right 1 – Go East – Turn Right 2 over nine days. We observed that hippocampal activity exhibited greater remapping between the first and second epochs of Turn Right from days 1-3 to days 7-9 in the One-Maze group mice, which performed the entire sequence on a single maze, compared to the Two-Maze group mice, which performed the Go East rule on a separate maze. Our results suggest that performing the Go East rule resulted in more remapping in One-Maze mice because they used a single representation of the spatial environment that was modified by performing the Go East behavior, while the Two-Maze group exhibited less remapping because they utilized a separate representation for the Go East rule. While both groups showed reinstatement of the Turn Right 1 representation during Turn Right 2, reinstatement was stronger in the Two-Maze group. We additionally showed that reorganization of neural activity affects the frame-by-frame coordination of activity among neural ensembles in part independently of remapping observed in session-averaged spatial event rate maps, similar to previous findings indicating this is a crucial, though less-studied, dimension of hippocampal activity (El-Gaby et al., 2021; Mau et al., 2022).

These findings are consistent with and expand on many previous studies. It is well known that hippocampal activity is often most strongly segregated by spatial environments, compared to other variables such as behavioral demand and the time interval between recordings, among many others (e.g. Leutgeb et al., 2005; McKenzie et al., 2014; Rubin et al., 2015). Our work directly compares the influences of spatial environment, changing behavioral task, and time interval on remapping. Previous work has established that new learning can cause changes in the place-related firing of hippocampal neurons (Alvernhe et al., 2011; McKenzie et al., 2013; Moita et al., 2004; Moita et al., 2003). Our study expands on these findings by fully reinstating the prior behavior (two epochs of Turn Right), using comparable tasks for the two behaviors (spatial navigation tasks), and by using calcium imaging to record neural activity over a long duration to expand the window over which these observations are made in the same neural population.

While many studies have measured remapping by comparing activity patterns before and after learning (Alvernhe et al., 2011; McKenzie et al., 2013; Mizuta et al., 2021), but without reinstating the behavioral conditions from the first task, it is impossible to exclude the possibility that differences in neural activity reported as changes are actually representations of each unique behavior or environmental configuration (Frank et al., 2000; Leutgeb et al., 2005; Markus et al., 1995; Mizuta et al., 2021; Wood et al., 2000). Memory updating has also been studied using across-group approaches (Zadbood et al., 2022). Previous studies have uncovered prospective effects on memory organization by using distinct training protocols in separate groups of animals before recording hippocampal activity from all animals performing the same task (Colgin et al., 2010; Plitt & Giocomo, 2021), and our study complements this work by examining retrospective effects by testing how new learning affects memories for prior experience. Our results are also consistent with results from the medial entorhinal cortex comparing free foraging before and after goal-directed navigation (Boccara et al., 2019; Butler et al., 2019).

Our task was designed to minimize the difference between the two behavioral demands by using two spatial navigation tasks with similar trial structures, start locations, trial durations and equal rewards. Fear conditioning and extinction studies also compare similar outward patterns of behavior before conditioning and after extinction, but it is possible the prior threat association of the environment constitutes a more fundamental change to the memory of that environment than does a different pattern of turn responses; outcomes with different valence relate to different survival instincts, which may engage fundamentally different memory processes (Chen et al., 2020). Even if animals do not display signs of threat expectation after extinction, the condition of the environment is changed in that safety is demonstrably not guaranteed (Zaki et al., 2021). The tasks used in our study may also engage different constellations of neural circuits, as previous work suggests that egocentric, body-turn responses (Turn Right) are supported by striatal circuits while allocentric responses (Go East) are supported by the hippocampus (Chang & Gold, 2003; McDonald & White, 1994; Mcintyre et al., 2003; Packard & McGaugh, 1996), although other studies have proposed that both tasks may be represented as arm-arm location associations in over-trained animals (Futter & Aggleton, 2006). We contend that two behavioral states described by different patterns of reward acquisition in a plus-maze are more comparable to each other than are the period following a shock and periods of free exploration before and after the threat of physical harm. Navigation tasks also allow for a longer window for observing activity when the hippocampus is known to be engaged than can be assumed during the window following a shock (Kentros et al., 2004; Pettit et al., 2022). However, as this study primarily concerns the degree of remapping observed across two behavioral epochs depending on whether an intervening behavior was performed in the same spatial environment, the results are consistent with the above-mentioned fear conditioning studies in that they demonstrate remapping following an intervening event, and additionally can be seen as expanding such findings to a novel combination of behaviors.

The behavioral paradigm in this study was also designed to address the separate influences of stabilization and orthogonalization on hippocampal representations. Memory stabilization begins immediately after a learning experience, continues for several days following, and includes consolidation (Alvarez & Squire, 1994; Kitamura et al., 2017) and changes to single neurons that refine their firing patterns and integrate them with sequences (van de Ven et al., 2016), among other processes. Hippocampal representations for experiences that have overlapping features, such as the spatial environment in which they occur, or happen close together in time are orthogonalized to prevent interference (McNaughton et al., 1991; Brown & Stern, 2014; Cai et al., 2016; Chanales et al., 2017; Gilbert & Kesner, 2002; Hasselmo & Eichenbaum, 2005; Hasselmo & Wyble, 1997; Law et al., 2016; McNaughton & Morris, 1987; O’Reilly & McClelland, 1994; Rashid et al., 2016). Orthogonalization appears as changes in the neural activity patterns causing them to become more distinct over time, and has been observed beginning early in the learning process (Cahusac et al., 1993; Chanales et al., 2017; Lee et al., 2006; Smith & Mizumori, 2006) and continuing for days after (Law et al., 2016; Lever et al., 2002). In these previous studies on orthogonalization, competing behavioral responses were performed on alternating trials or within minutes of each other, which means that subsequent changes to neural activity likely involve both orthogonalization and stabilization. Our study separates the onsets of the two experiences by several days, which should help to separate the effects of stabilization from orthogonalization. Additionally, previous studies suggest that memory updating manifests more strongly in neural representation following sleep-related consolidation (Speer et al., 2021). Our results additionally demonstrate that orthogonalization can reshape a memory representation first acquired several days in the past.

One result that remains to be explained is why greater change was observed in the One-Maze compared to Two-Maze mice on the South and East arms, where the behavior did not change, in addition to and in similar magnitude to the North and West arms where behavior did change. This finding is in contrast to other studies which have found changes to hippocampal activity specifically in the maze regions with divergent behaviors (Brown et al., 2010; Brown & Stern, 2014; Chanales et al., 2017; Lee et al., 2006; Smith & Mizumori, 2006). Piaget (1952) suggested that if new learning is sufficiently distinct from existing knowledge, then it can be “assimilated” into the memory network as a new trace without affecting pre-existing representations, and that when new learning overlaps with existing memory the existing representations will be “accommodated,” altered to integrate updated information with existing knowledge. One way that this framework has been codified is in the nonmonotonic plasticity hypothesis: weakly activated synapses are de-potentiated, strongly activated synapses are potentiated, and un-activated synapses remain unchanged (Newman & Norman, 2010). Support for this hypothesis has been found in physiological data (Hyman et al., 2003; Kwag & Paulsen, 2009; Newman & Norman, 2010; Pavlides et al., 1988) and in ensemble representations (Newman & Norman, 2010).

Interpreting hippocampal remapping studies using the non-monotonic plasticity hypothesis presents a challenge due to seemingly conflicting results. Some previous studies on remapping have found changes in hippocampal neural activity in only those firing fields associated with manipulated task features (Alvernhe et al., 2011; Cheng & Frank, 2008; Frank et al., 2004; Trouche et al., 2016), while others showed that manipulating a task feature can cause changes to place fields for distant locations and even for separate environments, an effect we term “distant remapping” (Gava et al., 2021; McKenzie et al., 2013). The nonmonotonic plasticity hypothesis suggests that only representations that are activated should be manipulated; if it is taken to be accurate, then, in the studies with distant remapping, those distant locations were in fact activated and altered. In the McKenzie et al. study (2013), while the new reward location was in a part of the maze away from the prior reward location, the animals were in the same spatial environment and they had to adjust their behavior to favor the new reward location over the prior one. The Gava et al. study (2021) demonstrated remapping in the familiar environment when animals learned a conditioned place preference in a separate pair of connected chambers, and remapping of the familiar environment was not observed when animals simply received a reward in a single-compartment novel environment. Neural activity that reflects changes in behavioral context has been modeled as changes in the forward sequence readout, including on short timescales where retrieval is gated by activity on the previous trial (Erdem & Hasselmo, 2012, 2014; Hasselmo & Eichenbaum, 2005); this mechanism could account for distant remapping that occurs within a spatial environment, as in the McKenzie study, but a broader view is required to account for distant remapping in separate spatial contexts.

We hypothesize that distant remapping occurs when there is competition between hippocampal representations for different behaviors within an environment. In the Gava study, reward in a novel environment with one compartment does not induce competition because there is no distinction to be learned; in the two-compartment conditioned place behavior, the mouse has to make a choice between compartments to visit in order to receive additional sucrose, leading to destabilization of the representation for the familiar context because it was experienced close in time even though it is physically separate. In the McKenzie study, the rat can receive rewards most efficiently by suppressing a visit to the prior reward location and stopping at the new reward location which it previously bypassed, meaning there is competition between old and new task parameters; the resulting destabilization can affect place fields around the track, suggesting that the representation for the track was a coherent structure in memory with respect to the behavior that could not be altered piecemeal. Distant remapping would not be expected in the Tolman detour task since the reward location is unchanged and choosing a less efficient route among open hallways does not stop the rat from reaching the reward (Alvernhe et al., 2011). In the study presented here, competition between representations for North-West and North-East responses in the same environment caused significant remapping on the South and East arms in One-Maze mice, even though behavior there was unchanged, presumably because those representations were closely related to representations for the North and West through close physical and temporal proximity since task occurred in a single environment with intermixed trials; competition may also have arisen to resolve the overlapping behavioral trajectories that end on the East arm. Meanwhile, Two-Maze mice were able to sleep between experiences of the two behaviors on different mazes, which may have reduced the temporal proximity between distinct behavioral demands enough to mitigate competition (like in Cai et al., 2016). In this perspective, competition causes widespread activation of representations that may be distant or remotely related in order to resolve representations for conflicting demands on behavior, invoking a wider kernel of non-monotonic plasticity than would be seen if each representation were recalled independently. To advance this hypothesis, future studies will need to explicitly delineate the conditions under which representational conflict occurs and the extent of the widespread activation and subsequent instability.

Previous work has demonstrated that the hippocampus is critical for using spatial context to disambiguate competing behavioral contingencies (Butterly et al., 2012), that changes to task demands can have impacts on established patterns of activity (Alvernhe et al., 2011; Boccara et al., 2019; Brown & Stern, 2014; Butler et al., 2019; Gava et al., 2021; McKenzie et al., 2013), and that the process of resolving competing demands on behavior can cause changes in representational organization (Brown & Stern, 2014; Chanales et al., 2017; Levy et al., 2021; Smith & Mizumori, 2006). We studied all these features in a single preparation over a long recording interval of days rather than minutes, which allowed us to characterize the impact of new learning on previously existing representations; to disentangle those effects from the interval between exposures and intervening changes in behavior; and to allow representations to begin to stabilize before introducing manipulations. Additionally, we use these results alongside previous literature to suggest a theoretical framework for considering the circumstances where remapping will be observed beyond the local range of a manipulation, offering new insight on the hippocampal memory system.

## Acknowledgements

This work was supported by the following grants: U.S. Office of Naval Research MURI N00014-16-1-2832 and N00014-19-1-2571, National Institutes of Health R01 MH052090, R01 MH 051570, MH060013, and MH120073. Thanks to Andy Alexander, Nat Kinsky and Will Mau for feedback and suggestions on the manuscript, to the entirety of the Hasselmo lab and many alumni for input and discussion, and to Dan Orlin, Wing Ning, Dan Sheehan, Jun Shen, Shelley Russek and Sandra-Jean Grasso for administrative support. The authors would like to acknowledge the GENIE Program, specifically Vivek Jayaraman, PhD, Douglas S. Kim, PhD, Loren L. Looger, PhD, Karel Svoboda, PhD from the GENIE Project, Janelia Research Campus, Howard Hughes Medical Institute, for providing the GCaMP6f virus. Thanks to Vardhan Dani and Laura Cardy at Inscopix for technical support with the imaging technology.

---

**Supplementary Figure 1:**
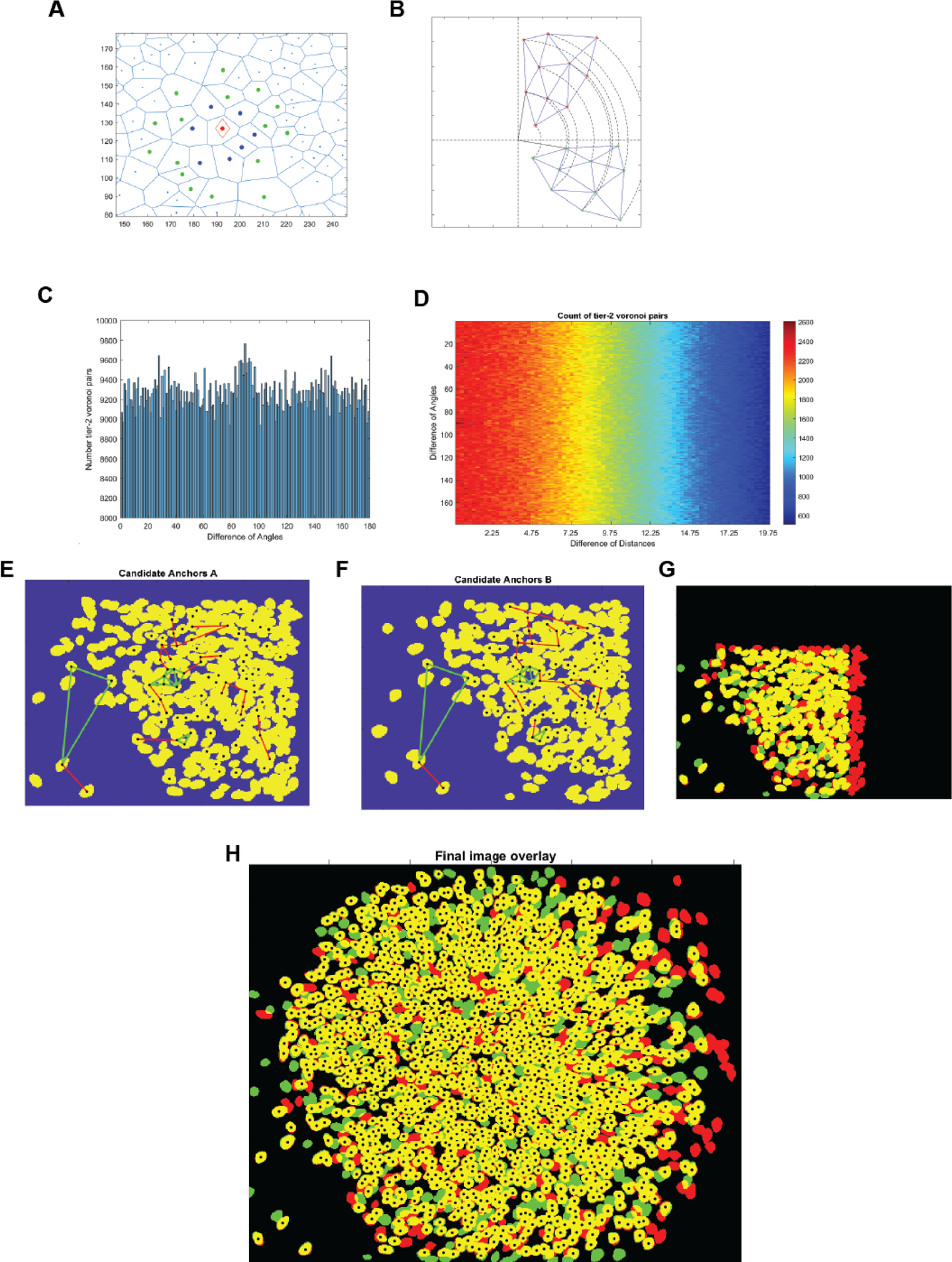
Diagram of steps for Voronoi-triangulation pre-alignment. **A**, Voronoi diagram showing a cell of interest (red diamond), Tier 1 adjacent points (blue) and Tier 2 adjacent points (green). **B**, example of the same set of points rotated 90 degrees. In the angle distance matrix for all points to all other points, correct alignment here would be a cluster at 90 degrees. **C**, Example angle differences histogram for real data, showing a peak near 90 degrees, registered FOV was intentionally rotated this far. **D**, 2D histogram for angle and distance differences; not the bin with the highest density at the smallest distance difference and 90 degrees offset. **E,F**, Black dots are potential anchor cells, lines are connected edges in the Voronoi diagram for this subset, and green lines are edges kept for satisfying quality matching criteria. E is the “base,” reference session, F the session to be registered. **G**, Alignment of these subsets of ROIs. **H**, Alignment of the whole field of view for both sessions.

**Supplementary Figure 2:**
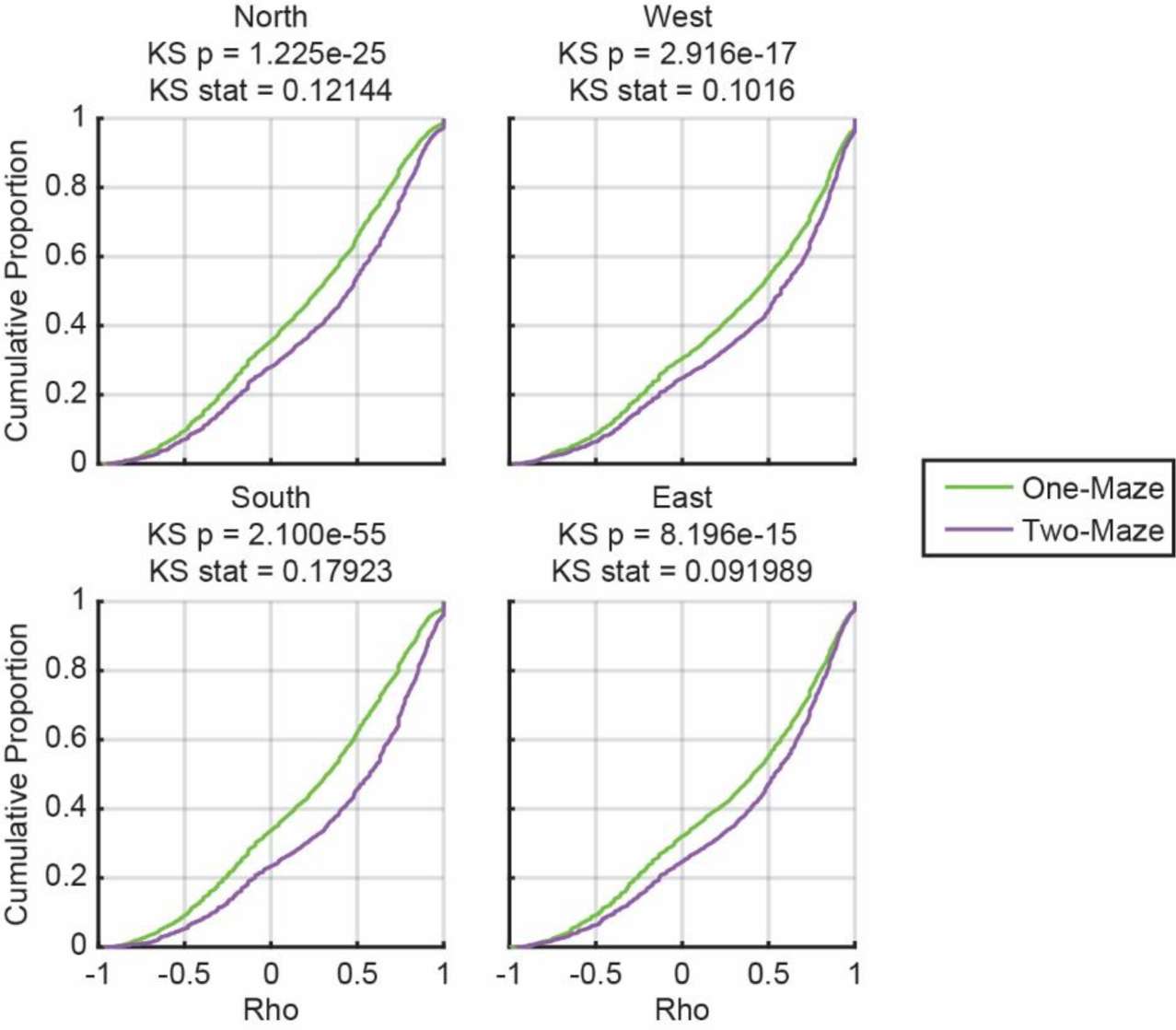
Single-Cell Event Map Correlations for Turn Right 1 – Turn Right 2 on Each Maze Arm. Same as in Figure 2, each plot shows the cumulative proportion of correlations of single cells from each pair of days in Turn Right 1 to Turn Right 2. Correlation calculated as Spearman’s Rho. Kolmogorov-Smirnov test results for each arm listed above each plot.

